# Neuronal components of evaluating the human origin of abstract shapes

**DOI:** 10.1101/085902

**Authors:** Hagar Goldberg, Yuval Hart, Avraham E Mayo, Uri Alon, Rafael Malach

**Affiliations:** Department of Neurobiology, Weizmann Institute of Science, Rehovot, Israel; School of Engineering and Applied Sciences, Harvard University, Cambridge, MA USA; Department of Molecular Cell Biology, Weizmann Institute of Science, Rehovot, Israel; The Theatre Lab, Weizmann Institute of Science, Rehovot, Israel

## Abstract

Communication through visual symbols is a key aspect of human culture. However, to what extent can people distinguish between human-origin and artificial symbols, and the neuronal mechanisms underlying this process are not clear. Using fMRI we contrasted brain activity during presentation of human-created abstract shapes and random-algorithm created shapes, both sharing similar low level features.

We found that participants correctly identified most shapes as *human* or *random*. The lateral occipital complex (LOC) was the main brain region showing preference to human-made shapes, independently of task. Furthermore, LOC activity was parametrically correlated to beauty and familiarity scores of the shapes (rated following the scan). Finally, a model classifier based only on LOC activity showed human level accuracy at discriminating between human-made and randomly-made shapes.

Our results highlight the sensitivity of the human brain to social and cultural cues, and point to high-order object areas as central nodes underlying this capacity.

## Introduction

Humans are social and tuned to social cues. They therefore need to distinguish true social cues from other stimuli. Previous studies have shown that human observers can distinguish biological motion from random motion, even in impoverished stimuli such as point-light displays (Johansson, 1973). This is suggested to be an intrinsic ability of the visual system, which has evolved to preferentially attend to other humans, as shown in newborn babies (Simion et al., 2008). These findings suggest that essential information such as social recognition can be derived from minimalistic dynamic displays. In addition to direct communication (through verbal and body language) human culture has ways of indirect communication through visual symbols - a conventional representation of concepts through scripts and art; starting from cave paintings (Chauvet et al., 1996), and continuing nowadays with symbols such as emoticons.

Similarly to the way humans can distinguish human walk from random motion generated by spot light displays, we hypothesize that humans can successfully recognize abstract shapes that have been generated by other humans compared to similar shapes created randomly. We further hypothesize that the neuronal mechanisms that underlie this capacity are likely to be ingrained in core systems and hence independent of the task performed.

What could be the aspects of the shapes that underlie the ability of participants to categorize them into *human* vs. *random*? The symbolic meaning of a shape and its familiarity can serve to assess the shape’s origin. Another relevant feature is the shape’s aesthetic value, if indeed people tend to create more beautiful shapes. Thus, in the present study we focused on beauty and familiarity- i.e. iconicity - the sense that a shape represents a familiar icon. We also examined the inverse of familiarity- i.e. “weirdness”- a subjective sense that a figure is strange and unfamiliar. Both beauty and weirdness are intuitive and powerful yet subjective impressions which are difficult to define. Substantial work has suggested beauty and weirdness carry evolutionary advantages. For example, neuro-imaging study showed overlap in brain regions which function both during processing of aesthetic artworks and during appraisal of evolutionary important objects (e.g. attractive potential mates, or desirable food) (Brown et al., 2011). Visual artistic representations of beauty (Kawabata and Zeki, 2004, Di Dio et al., 2007, Ishizu and Zeki, 2011, Cattaneo et al., 2015, Vartanian and Goel, 2004, Lacey et al., 2011) or naturalistic stimuli beyond the arts domain (Brown et al., 2011, Chatterjee et al., 2009, Kirk, 2008, Lacey et al., 2011) were showed to be linked to experience of reward, pleasure, and attitudes to external information (approach/withdrawal). Similarly, the concept of weirdness was attributed to signals of danger and risk (Rotshtein et al., 2001). Sensing beauty and weirdness thus seem important for human behavior and survival, but we have little understanding of their neuronal correlates.

Another line of work considered the contrast between artificial and familiar images in a number of studies of the human visual cortex (e.g. Fourier descriptors, and scrambled images; Lerner et al., 2002, Tsao et al., 2003, Aalto et al., 2002, Murray et al., 2002, Malach et al., 1995). Visual objects such as face images were rendered bizarre by inverting internal face features (Rotshtein et al., 2001). These studies showed an increased activity of high order object areas (lateral occipital complex - LOC) to coherent objects whether they are familiar or unfamiliar (Malach et al., 1995, Kanwisher et al., 1997, Kanwisher et al., 1996, Grill-Spector et al., 2001, Rotshtein et al., 2001).

However, in previous work a control over low-level features was lacking. In fact, we are not aware of systematic study that directly compared brain activation to human-generated vs. randomly generated shapes constructed of similar low-level components. Furthermore the contribution of beauty and familiarity to the ability to evaluate human origin of shapes has not been explored directly in previous research.

To address these issues, we examined, using fMRI, human perception of simple shapes made by other humans, compared to similar shapes generated by an artificial algorithm. We specifically examined to what extent brain regions respond differentially to these two categories - and to what extent the beauty, iconicity and weirdness dimensions contribute to this differentiation. To this end we employed a novel design in which a large group of 101 people were asked to create simple shapes and to rank them according to how appealing they were (interesting and beautiful) (Noy et al., 2012). Additional shapes were generated by an artificial, random walk algorithm sampling from the space of all possible shapes and excluding shapes that were generated by human observers. The generated shape ensemble included a wide spectrum of beauty and familiarity levels.

We found that participants which were unfamiliar with the shapes were able to successfully distinguish between human and random origin shapes. Our brain imaging results show that LOC activity was significantly higher for the human-made shapes compared to the *random* ones. Furthermore, this differential activity was a combined result of a positive correlation to beauty and iconicity (familiarity) and a negative correlation to weirdness (unfamiliarity) of the abstract shapes regardless of task. Our results point to the high-order object related complex (LOC - Malach et al., 1995) as a pivotal node in endowing human observers with the ability to recognize shared symbolic meaning and distinguish human from artificially created shapes.

## Results

Here we aimed to study the behavioral and neuronal mechanisms of distinguishing whether an abstract shape was created by a human from a given space of shapes or by an algorithm that makes a random choice from the same space of shapes. We used a rich yet fully determined space of shapes, made of ten contiguous squares. Shapes were built by either 101 human players in a computer-shape-generation game (see methods and Noy et al., 2012) or a random choice algorithm from the same space of shapes. Thus, we contrasted algorithm-created with human-created abstract shapes having similar low-level features. The human-created shapes were further subdivided into three groups of different appeal ratings (by their human creators, see Methods). Subjects unfamiliar with the shapes underwent fMRI scanning while watching the shapes in a block design. The shapes were presented in two different experiments. In the first, subjects performed a color discrimination task (experiment and task 1), and in the second, a human vs. random algorithm origin discrimination task (experiment and task 2). Following the scan, subjects gave their subjective evaluations of beauty, weirdness and iconicity of the shapes (see Methods for details).

### Behavioral results

Subjects successfully distinguished between most human and random creation shapes (Fig. 2a). Most categories were successfully classified by the subjects (85-90% accuracy), except *not chosen* category which was correctly classified (as *human*) in only 30% of the cases (chance level = 50%). This difference in success rate between *not chosen* category and the other categories was significantly lower (Mann-Whitney U = 0, n_1_ = n_2_ = 7 p < 0.005, twotailed). The difference between all other categories was not significant. A bias-free signal detection analysis indicated that subjects were able to reliably distinguish between human and random made shapes (N = 13, mean d’ = 2.03, SE = ±0.34). No difference in reaction times was found between the four categories in both experiments. Three subjective aspects of the shapes were examined; beauty, weirdness and iconicity. Weirdness and iconicity scores complemented each other – while weirdness focused mainly on the level of unfamiliarity of a shape, iconicity reflected the level of familiarity and distinct meaning. The classification to human origin was best modeled by the interaction of the beauty and weirdness scores rather than the two scores separated (*P*(*human*) = (1 + 13.7 *e*^−6.6 *WB*^)^−1^) (see Methods and SI). Beauty scores were lowest for *random* and *not chosen* shapes, and were significantly higher for *chosen* and even higher for *top rate* (Mann-Whitney between *random* and *not chosen*: U = 20, n_1_ = n_2_ = 7, p = 0.521, two-tailed; between all other categories: U = 0, n_1_ = n_2_ = 7, p < 0.005, two-tailed. see Fig. 2b). Thus indicating cross population consistency between the creators of the shapes and the scanned participants.

**Figure 1.**
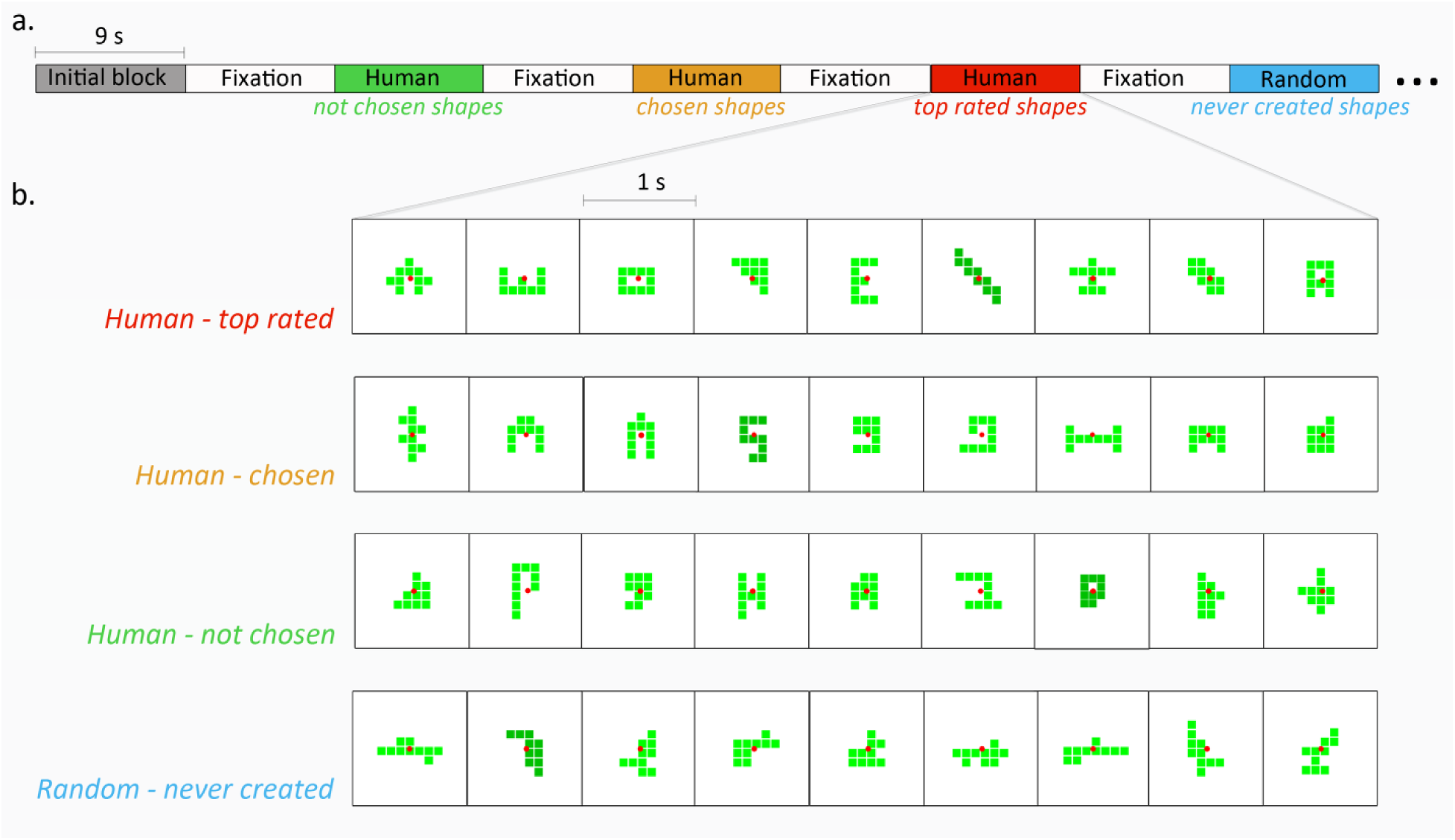
Experimental design. (a) Experimental protocol: shape images from four categories were presented in a block design with 9 second blocks, followed by 9 second fixation screen. Shape categorization was based on its creation process during a computer game in a previous work (Noy et al., 2012, see methods); *Top rated shapes* were created by human players and rated as most creative shapes. *Chosen shapes* were created by human players, chosen as beautiful shapes but never rated as most creative shapes. N*ot chosen shapes* were created by human players but never chosen as beautiful shapes or rated (by the players). *Never created shapes* were created by a random algorithm, and never by human players. (b) Example for a block of each category. Each block included 9 images (one second each) of the same category; 8 images in light green and one in dark green.

**Figure 2.**
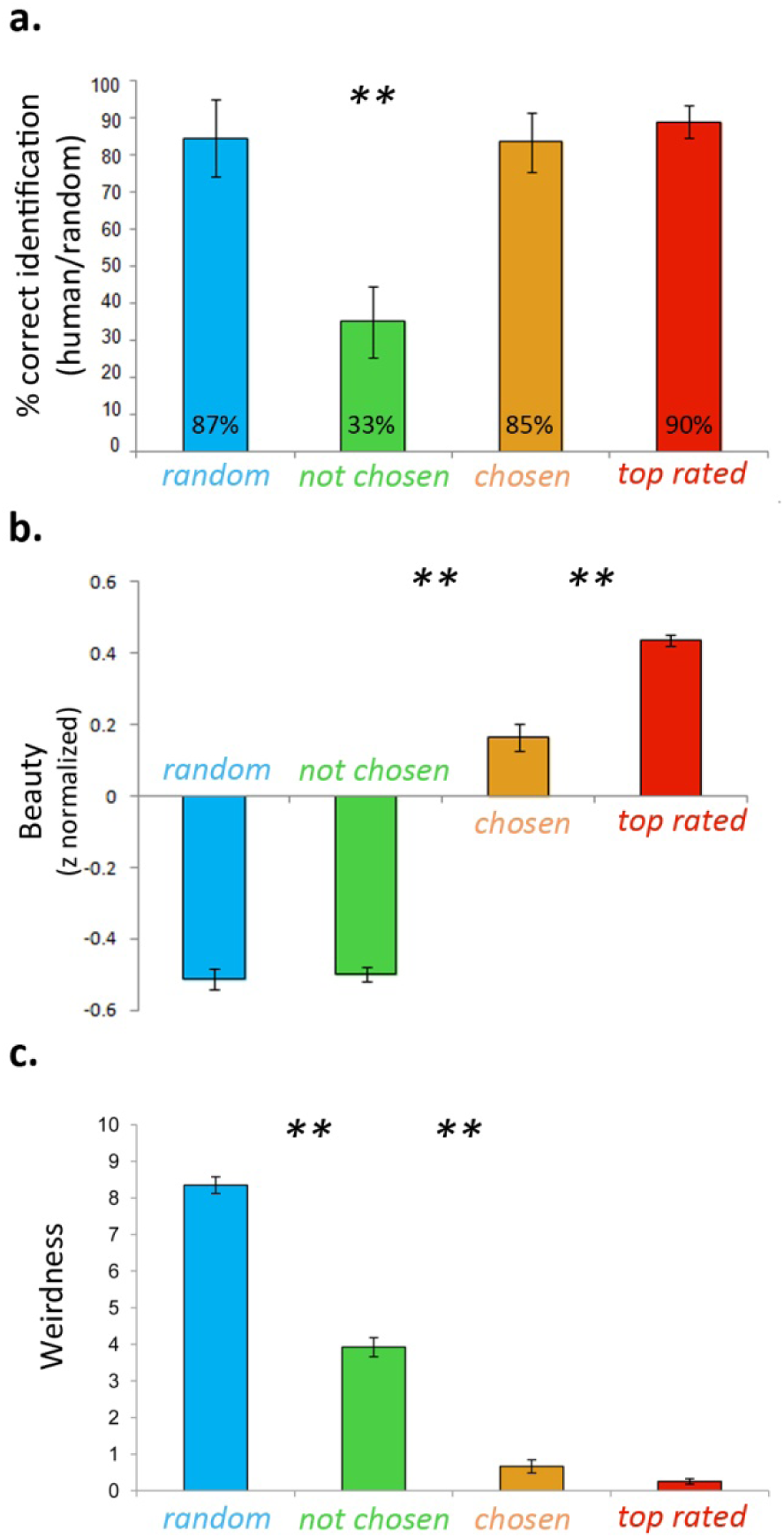
Behavioral measurements. (a) Group’s average percent success rate (human/random) in each category; *Human- top rated, Human- chosen, Human- not chosen, Algorithm-random*. Two-tailed, within subjects Mann-Whitney between categories, n1 = n2 = 7, p < 0.005**, error bars indicate the groups’ standard error. (b) Group’s average beauty scores per category (z-score normalized). (c) Group’s average weirdness scores per category.

Weirdness scores on the other hand distinguished between *human* and *random* categories; the *random* category received the highest score, *not chosen* received a significantly lower score, *chosen* and *top rated* received the lowest scores, significantly different from the first two (Mann-Whitney between *chosen* and *top rated*: U = 0.531, n_1_ = n_2_ = 7, p = 0.08, two-tailed; between all other categories: U = 1, n_1_ = n_2_ = 7, p < 0.005, two-tailed. see Fig. 2c). Thus, subjective beauty was positively correlated, and weirdness was negatively correlated to classification as *human*. Furthermore each parameter separated between different shape categories.

### Brain imaging results

In order to examine a possible implicit differentiation between human-made shapes and randomly made shapes, a direct contrast of BOLD activity (task 1) between *human* blocks and *random* blocks was conducted. The contrast map (Fig. 3) revealed highly localized preferential activations to *human* vs. *random* in the lateral occipital complex (LOC, Malach et al., 1995). Preferential activation to *random* shapes was spread over a wider range of the cortex, particularly in parietal and frontal regions. Its most significant activation was located in the inferior parietal gyrus (IFG).

**Figure 3.**
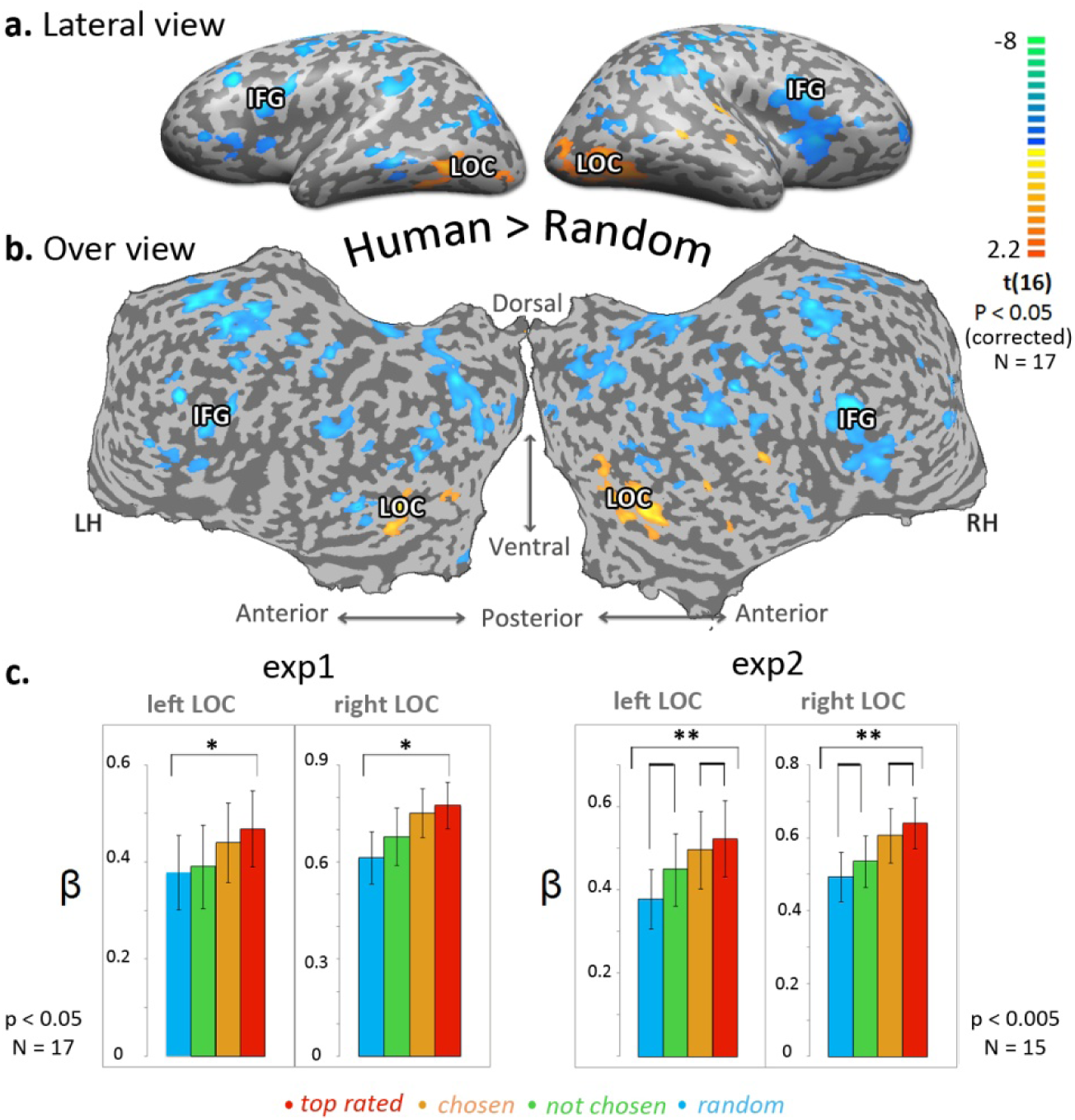
Comparison of Human and Random cortical activations. All human generated shapes *(Human)* vs. computer generated shapes *(Random)* multi subjects activity map (experiment 1, N = 17, corrected p < 0.05) is presented on an inflated cortex, in a lateral view (a) and unfolded cortex (b). Color scale indicates t values. Yellow-orange scale represents regions which were more activated while watching human generated blocks compared to random shapes (blue-green scale). Most significant activated regions are marked on the inflated map; Lateral Occipital cortex (LOC), Inferior frontal gyrus (IFG). (c) LOC ROI analysis: repeated measures ANOVA between averaged beta values of each category; *Human- top rated, Human- chosen, Human- not chosen, Algorithm-random*. Left side - experiment 1 (N = 17, p < 0.05*), right side - experiment 2 (N = 15, p < 0.005**).

Cross-task ROI analysis demonstrated a consistently stronger response to *human* blocks relative to *random* blocks in bilateral LOC in both tasks (paired t test; (experiment 1) left LOC, N = 17, t = 2.74, p < 0.05; right LOC, N = 17, t = 3.92, p < 0.005; (experiment 2) left LOC, N = 15, t = 33.93, p < 0.005; right LOC, N = 15, t = 3.29, p < 0.05). ROI analysis inspecting the averaged LOC activity (beta weight) per category revealed a gradual positive response in bilateral LOC along the “appeal” axis (Fig 3c). *Random* shapes showed the lowest response in LOC, top rated showed the highest response (followed by not chosen and chosen respectively) (One way repeated measures ANOVA; (experiment 1) left LOC, N = 17, F = 2.751 not significant, Linear trend: F = 5.174, p < 0.05; right LOC, N = 17, F = 4.35, p < 0.05, Linear trend: F = 7.974, p < 0.05; (experiment 2) left LOC, N = 15, F = 6.145, p < 0.05, Linear trend: F = 28.721, p < 0.0005; right LOC, N = 15, F = 8.052, p < 0.0005, Linear trend: F = 26.281, p < 0.0005).

Since beauty, weirdness and iconicity scores were predictive for blocks classification (*human/random*) we wanted to study their neuronal correlates. A whole brain parametric GLM analysis was therefore conducted (Fig. 4). The results revealed a consistent parametric relationship between LOC activity to beauty, weirdness and iconicity measures regardless of task (color/ shape origin discrimination). Parametric brain maps for beauty scores show focused activation in the LOC. However in frontal areas, weirdness and iconicity showed a task-related selectivity: in task 1 frontal region such as dorsal premotor cortex and inferior frontal gyrus (IFG) were parametrically correlated to weirdness (positive) and iconicity (negative). In task 2 lateral frontal cortex and superior parietal lobe showed parametric correlation to both weirdness (negative) and iconicity (positive) measures (Fig. 4). Finding a common neuronal network to both weirdness and iconicity with inverse correlation, supports the idea that these parameters reflect two opposite aspects of shapes familiarity and symbolic meaning.

**Figure 4.**
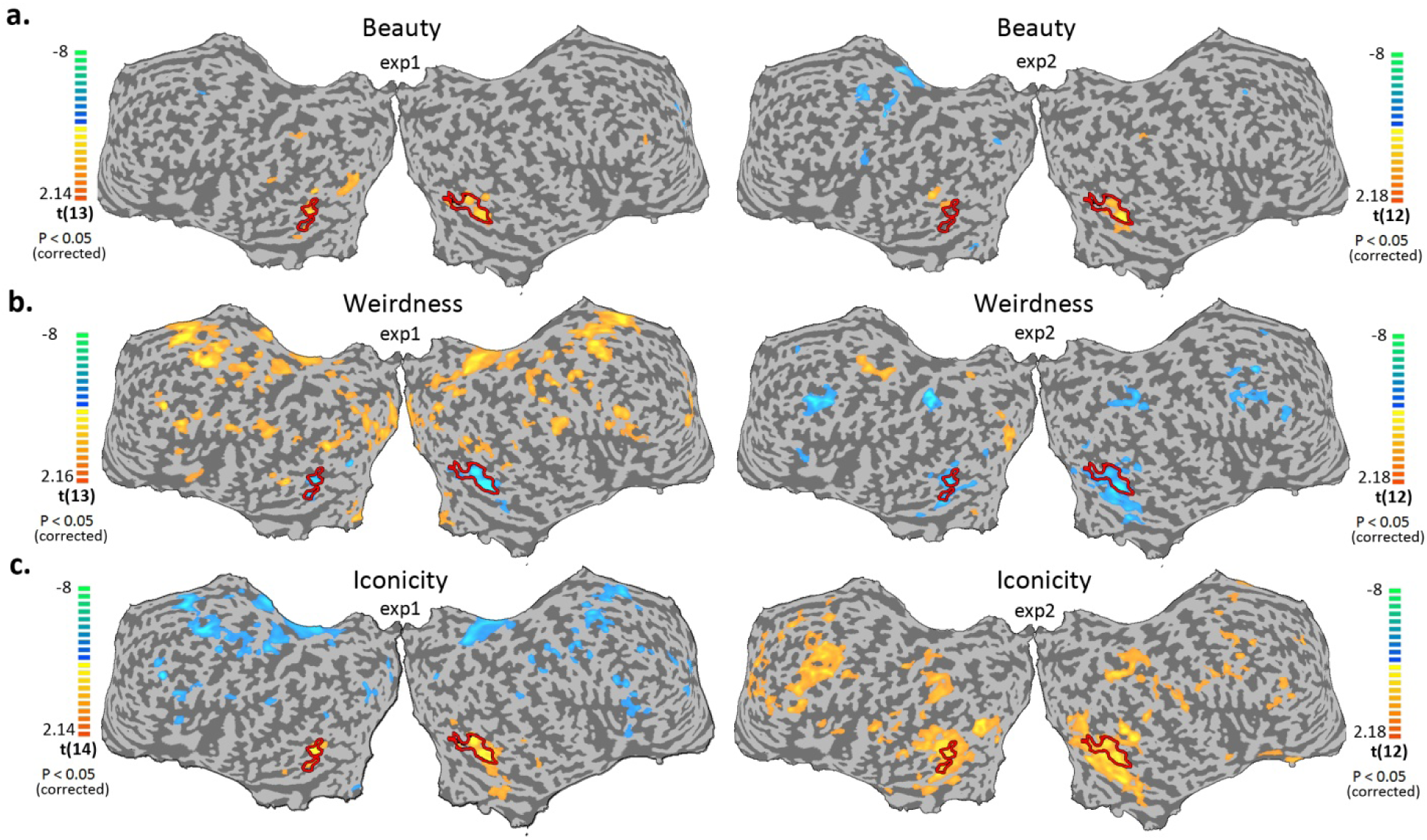
Parametric mapping. Cortical activity maps of multi subjects, random effect, parametric GLM analysis of beauty (a), weirdness (b), and iconicity (c) presented on an unfolded cortex, right side experiment 1 (N = 13, 13, 14), bottom panel experiment 2 (N = 12, 12, 13), both maps are corrected for multiple comparisons, p < 0.05. Color scale indicates t values. Yellow-orange scale represents regions which showed positive parametric relation with beauty/weirdness/iconicity scores. Blue-green scale represents regions which showed negative parametric relation with beauty/weirdness/ iconicity scores.

In order to disentangle the coupling between the subjective parameter of beauty and the objective parameter of symmetry (Spearman correlation, r = 0.92, p < 5*10^−11^), we calculated a subjective beauty score – blocks were ranked by their beauty score per subject, and then beauty scores of blocks with same ranking were averaged across subjects. The result is a perceptual beauty score which is independent of physical attributes such as symmetry. ROI analysis demonstrated a significant positive correlation between both symmetry/beauty and LOC activity (averaged normalized beta weights). However, the subjective beauty scores showed a significantly higher correlation compared to the symmetry scores (Spearman correlation; (experiment 1) left LOC, Symmetry: r = 0.4, ns; Beauty: r = 0.5, p < 0.005; right LOC, Symmetry: r = 0.47, p < 0.05; Beauty: r = 0.72, p < 5*10^−5^; (experiment 2) left LOC, Symmetry: r = 0.57, p < 0.005; Beauty: r = 0.69, p < 5*10^−5^; right LOC, Symmetry: r = 0.5, p < 0.05; Beauty: r = 0.81, p < 5*10^−7^, see Fig. 5). The correlation scores were further compared using bootstrapping method, where data points were randomly chosen (with replacements) from each dataset and the correlation was calculated. Each process was repeated 1000 times. (Spearman correlation; (experiment 1) left LOC, Symmetry: r = 0.28 ±0.17, Beauty: r = 0.54 ±0.13, d’=1.2; right LOC, Symmetry: r = 0.18 ±0.17, Beauty: r = 0.72 ±0.18, d’=2.9; (experiment 2) left LOC, Symmetry: r = 0.35 ±0.15, Beauty: r = 0.69 ±0.06, d’=2.1; right LOC, Symmetry: r = 0.28 ±0.16, Beauty: r = 0.81 ±0.05, d’=3.1. All values are Mean ±STD, N_(symmetry)_ = 16, N_(beauty)_ = 13). According to these results perceptual beauty showed a graded response across the entire dynamic range enhancing correlation and significance while symmetry correlation highly depended on the points chosen as evident from its large correlation variation (Fig. 5).

**Figure 5.**
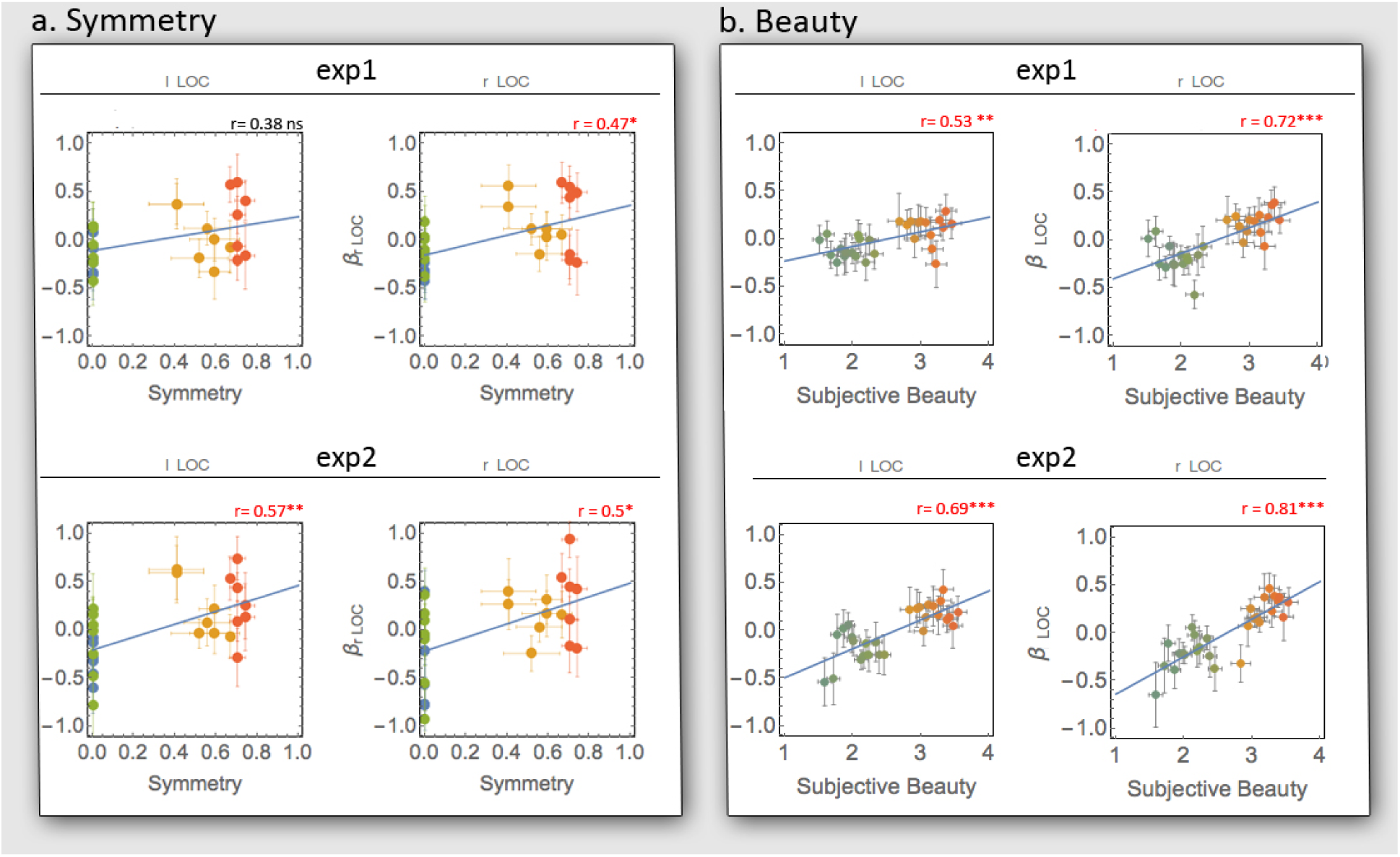
Correlations of LOC activity and symmetry/beauty. (a) Scatter plots present the relation between averaged blocks’ symmetry (x axis) and group’s averaged brain activity (normalized beta weight, y axis) in bilateral LOC. Each dot represents one block, and color indicates the blocks’ category (*Top rated* in red, *chosen* in orange, *not chosen* in green, *random* in blue) N = 13. Spearman correlation; (experiment 1) left LOC, r = 0.4, ns; right LOC, r = 0.5, p < 0.05. (experiment 2) left LOC, r = 0.6, p < 0.005; right LOC, r = 0.5, p < 0.05. (b) The relation between group’s averaged beauty ratings (x axis), and the group’s averaged brain activity (normalized beta weight, y axis) in LOC. Each dot represents mixed blocks with similar beauty ranking, color indicates the blocks’ category. N=16. Spearman correlation; (experiment 1) left LOC, r = 0.52, p < 0.005; right LOC, r = 0.69, p < 0.0005. (experiment 2) left LOC, r = 0.69, p < 0.0005; right LOC, r = 0.83, p < 0.0005.

Iconicity group scores were also correlated to LOC activity; Spearman correlation; (experiment 1) left LOC, r = 0.51, p < 0.01; right LOC, r = 0.65, p < 0.0005. (experiment 2) left LOC, r = 0.65, p < 0.0005; right LOC, r = 0.57, p < 0.005.

Lastly, we tested whether one could infer a shape’s origin by the beta activity of LOC. We compared the average LOC activity for each block with its probability to be classified as human. We fitted a non-linear hyperbolic tangent classifier with a random partial sample of data points (23/28) and iterated the process 1000 times. We found that the averaged fitted function predicted 78% of the shapes accurately by using the average beta activity of the block (compared to 50% for chance performance). Interestingly, the classifier’s ability resembled the subject’s behavioral results with 86% correct in *random*, *chosen* and *top-rated* categories, and only 57% in the *not-chosen* category. Thus, LOC activity might serve as a predictor to the shape’s origin.

## Discussion

The human brain is skilled in distinguishing between the familiar and the strange, between the natural and the artificial, and here we examined its ability to distinguish between human and random creations. We showed here a tight connection between behavior and brain function related to the process of identifying the origin of human-generated versus random abstract shapes. Our findings revealed that human subjects could correctly identify shapes as human made or randomly made. Subjects showed a general agreement that *random* shapes appeared weirder. Moreover their aesthetic evaluation of the shapes (Fig. 2b) was very similar to the evaluation made by the shapes’ creators (the three categories of human made shapes - see methods). These findings suggest a general agreement among individuals about what is considered human, meaningful and beautiful.

While previous studies used complex naturalistic images or objects, in the current study we used relatively simple and well-controlled shapes. Although similar in low level features, some of them (*random* shapes) were out of the common human scheme as manifested by the players playing the game. Indeed they were never created by human players (although probabilistically they should have been created), and they were perceived differently by the human brain.

Brain activity - specifically, the LOC, showed preferential activation to *human* relative to *random* blocks even in task 1 (color discrimination) in which attention was targeted to color rather than shapes. This supports our hypothesis that the human brain is capable of recognizing human creation even when not explicitly instructed to do so.

The LOC, well established as a hub of visual object recognition (Malach et al., 1995, Grill-Spector et al., 2001) - was the central region showing a preferred activation to human-made shapes compared to *random* shapes (task independent). The complementary, preferred activation to *random* shapes was found in parietal regions, motor cortex and most significantly in the inferior frontal gyrus (IFG, see figure 3). These findings are compatible with previous studies which reported that IFG is responsive to unexpected stimuli (Huettel and McCarthy, 2004), and to incongruent stimuli specifically within a social context (Shibata et al., 2011).

We examined 3 subjective shape characteristics with predictive value to classify as *human/random;* beauty, weirdness and iconicity. Using a whole brain analysis exploratory approach - we searched brain networks which were parametrically connected to each external measure. These analyses (Fig. 4) showed that LOC was a central node - parametrically correlated to shapes’ beauty, weirdness (inversely correlated) and iconicity. Moreover, using LOC activity might allow predicting shape’s origin with human level accuracy (Fig. 6). Thus, our findings point to the LOC as a central node for human vs. random origin, with shape’s beauty and symbolic meaning playing a role in that evaluation.

**Figure 6.**
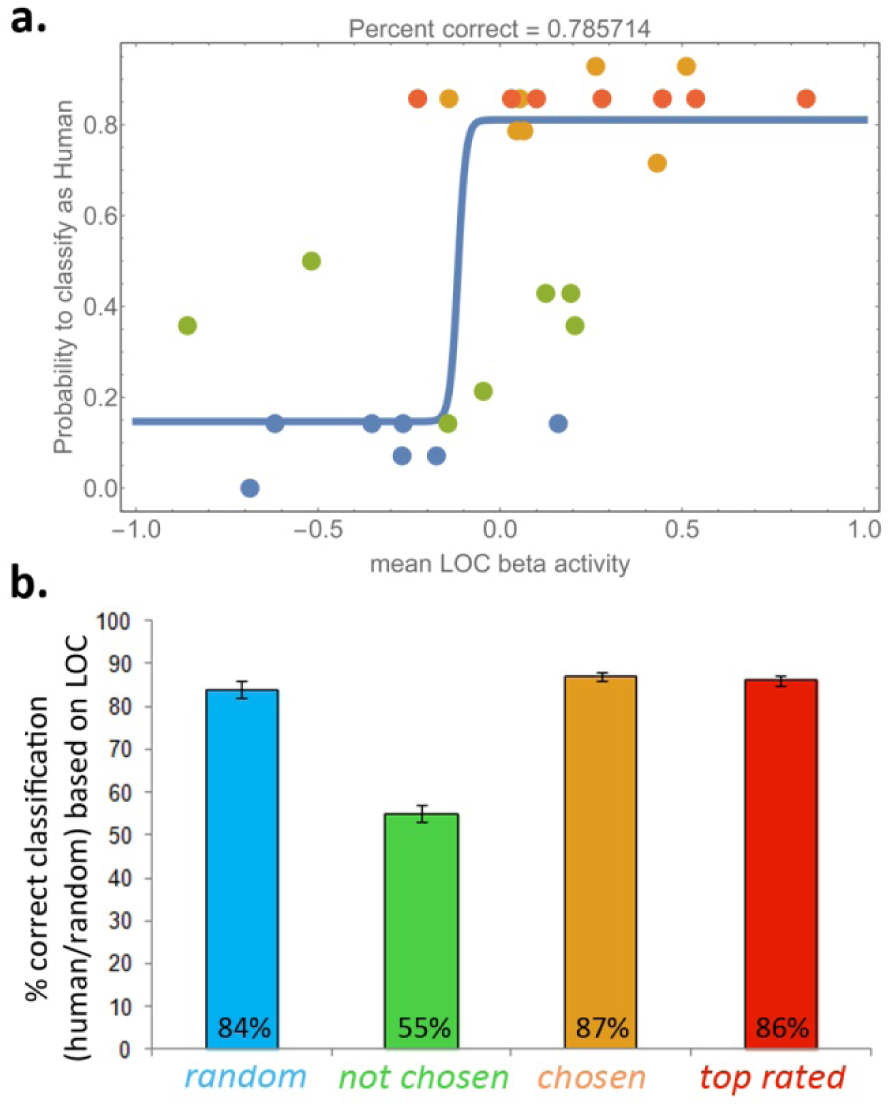
Classification of blocks’ origin based on their average LOC activity results in human-level accuracy. (a) We fitted a hyperbolic tangent classifier, *p*(*human*) = *a + b Tanh[b x + d]* (where x is taken to be the averaged LOC activity of the block) to the data points by a bootstrapping method. Each iteration, 23/28 points were chosen randomly and the best fit parameters were extracted. We repeated this process 1000 times and averaged the parameters of all the runs. Shown is the averaged model. Model parameters are (mean±ste): a=0.48±0.03, b=45±9, c=5±1, d=0.33±0.04. (b) Classification accuracy of the classifier on the different categories (top rated, chosen, not-chosen, random). Error bars are STE and are calculated by 100 random sampling with replacements of the real data points per each category.

### Brain and beauty

Previous studies of beauty, focusing on response to visual arts or objects perception, found activations in reward related areas, such as the orbito-frontal cortex (Kawabata and Zeki, 2004, Brown et al., 2011, Ishizu and Zeki, 2011, Kirk, 2008, Lacey et al., 2011, Cela-Conde et al., 2004), in emotion related areas like the amygdala (Brown et al., 2011, Di Dio et al., 2007, Ishizu and Zeki, 2011) and insula (Brown et al., 2011, Di Dio et al., 2007) and also in motor areas (Kawabata and Zeki, 2004, Ishizu and Zeki, 2011), pointing to a link between the experience of beauty to reward or information gathering and response upon it.

Recent studies also point to high order visual areas, and specifically the LOC, as an aesthetic center in the brain (Cattaneo et al., 2015, Chatterjee et al., 2009, Kirk, 2008, Lacey et al., 2011, Vartanian and Goel, 2004). Some reported that high order visual areas show preferential activity for aesthetic value only for representational art and not for abstract art (Cattaneo et al., 2015, Vartanian and Goel, 2004).

In the current study we explored which brain regions were sensitive to the beauty of abstract shapes. Our results revealed a consistent (experiments 1 and 2) positive parametric correlation between subjective beauty scores and bilateral LOC (Fig. 4a). The consistency and specificity of LOC in beauty analyses, suggests that beauty evaluation of abstract shapes is an automatic - bottom up process which is less dependent on the attention to the shape.

Previous studies pointed to the LOC as a central player in symmetry evaluation (Hodgson, 2009, Chatterjee et al., 2009, Beck et al., 2005, Sasaki et al., 2005). In the present study symmetry indeed showed a significant correlation to beauty ratings. Importantly, the correlation between LOC activity and symmetry was significantly weaker and less reliable compared to the beauty score correlations (Fig. 5). Thus, our paradigm of abstract shapes allowed the decoupling of the symmetry measure from that of the beauty scores. Our results show that subjective beauty evaluation is better correlated to LOC activity compared to shapes’ symmetry.

### Brain and familiarity

According to our results - both weirdness and iconicity, measures of familiarity and symbolic meaning, were correlated with LOC activation. Previous works showed increased activation in LOC which was similar for familiar and unfamiliar objects as long as the object is a coherent one (Malach et al., 1995, Kanwisher et al., 1997, Kanwisher et al., 1996, Grill-Spector et al., 2001), tracing a difference in activation between coherent objects response to non-coherent ones (using Fourier descriptors, and scrambled images; Lerner et al., 2002, Tsao et al., 2003, Aalto et al., 2002, Murray et al., 2002, Malach et al., 1995). This is compatible with the earlier finding of an enhanced activity in mid-fusiform gyrus in a response to abstract shapes, associated with long-term familiarization (Gauthier et al., 1999). Similarly, right LOC showed enhanced activity to abstract object structures following a short-term learning (familiarization) (de Beeck et al., 2006). Furthermore, LOC preferential activity to familiar (iconic) objects was shown in a visual imagery with haptic perception task (Deshpande et al., 2010, Lacey et al., 2010).

Our abstract shapes paradigm introduced a broad range of familiarity levels, which allowed to explore the effect of familiarity on LOC activity on a graded scale instead of a binary approach (familiar/unfamiliar) and without distortion of object’s features (Lerner et al., 2002, Tsao et al., 2003, Aalto et al., 2002, Murray et al., 2002, Malach et al., 1995, Rotshtein et al., 2001). Shapes with different familiarity levels manifested a difference in LOC activation although composed of similar low level features.

Interestingly, weirdness and iconicity showed inversed correlation patterns within very similar neuronal networks, supporting the idea that they reflect the two ends of shapes familiarity and meaning (not only theoretically but also neuronaly). While the beauty parametric analysis showed a consistent and local activity of LOC, both weirdness and iconicity parametric mapping showed a task dependent activation, and a broader network involvement, including frontal and parietal regions in addition to LOC (Fig. 4b,c).

It could have been argued that the identification of human origin shapes was mainly due to highly recognizable symbols (e.g. letters and digits). However- such iconic shapes were only a small portion (~10%) of the entire ensemble (sup Fig. 1). In fact, the shape ensemble introduced a gradient of iconicity as was shown by both ratings and brain responses. Furthermore, it has been well established that letters and digit representations are left- lateralized (Hasson et al., 2002, Fiez and Petersen, 1998, McCandliss et al., 2003) while we consistently found a slight right-hemisphere bias of the *human* vs. *random* contrast (sup Fig. 2).

De Beeck et al have showed an overall increased response to trained abstract objects compared to untrained abstract objects. Right LOC in particular showed the strongest increased response as a result to the training (de Beeck et al., 2006). In our study there was no formal training and all the shapes were novel, however we suggest that the human schema is an expression of an inherent training for social representations in the human mind. This schema led human players to explore certain shapes, and avoid other shapes (unlike the random walk algorithm), similarly this schema guided participants to successfully distinguish between human made and randomly made shapes. It is an open question whether this schema was developed as a result or is the source of human graphic communication. An interesting future avenue related to this question will be to investigate whether these findings are reproducible across cultures, within cultures, and in young infants. It will also be of interest to study whether humans on the autistic spectrum experience and neuronaly process these shapes similarly to typical individuals.

### Conclusion

Our study examined the behavioral and neuronal components of evaluating human origin of abstract shapes. Our results point to high order object areas (LOC) as a central node in beauty representation, and in symbolic meaning attribution of abstract shapes. Both aspects had a significant contribution to the classification of shape origin as *human* or *random*, and may have a key role in visual human communication (such as visual art).

## Materials and methods

### participants

Seventeen healthy right handed subjects (ages 28 ± 3.8, 10 females) participated in the fMRI experiments. Fifteen of them participated in both experiments 1 and 2. Fourteen filled a beauty, iconicity and weirdness evaluation questionnaire post the fMRI scan.

### Task and stimuli

Shape stimuli. Shapes of ten contiguous identical green squares were created in a shape-search computer game, by either 101 human players or a random walk algorithm (Noy et al., 2012). In each shape, squares were connected by an edge. Players moved one square at a time to create new shapes. They were instructed to place beautiful and interesting shapes into a gallery by pressing a button. There was no limit to the gallery. Players played for 15 min and created about 310 shapes, of which they chose 46 shapes on average to the gallery. At the end of the game, players chose the 5 most creative shapes from their own gallery. The shape space includes 36,446 possible shapes.

Shapes were classified into four categories based on their origin (human/algorithm), and their appeal ratings (by their human creators). *Not chosen shapes* were created by human players but never chosen as beautiful shapes (by the players). *Chosen shapes* were created by human players, chosen as beautiful and interesting to the gallery but never rated as most creative shapes. *Top rated shapes* were created by human players, chosen as beautiful and interesting shapes and were rated by players as most creative shapes. *Random* (*Never human-created shapes)* were created only by a random walk algorithm (and never by human players) on the space of shapes, where the next shape is chosen randomly from neighboring shapes that are one move of a square away from the current shape. Length of walks was sampled from the distribution of walk lengths of the human players. For each category, the 20 most frequent shapes, i.e. the shapes that were the most common to many players (or random walks in the random category), were chosen for the fMRI experiment (see the shapes in sup. Fig. 1).

### Experimental design

During the fMRI scan the created shapes were presented in homogeneous-category blocks lasting 9 sec, followed by a 9 sec fixation screen. Each block consisted of 9 images (one second each); eight images in light green and one image in dark green. Each category was presented in seven different blocks. To reduce scan novelty effect, an extra block (which was not analyzed) was added to the beginning of each experiment, 29 blocks were presented in total.

Each subject watched the same sequence of blocks twice (once for each task). In the first experiment (task 1) subjects were required to classify the stimuli according to color; light green (press 1) or dark green (press 2). In the second experiment (task 2) following each block, subjects were required to classify the shapes of the preceding block as human creation (press 1) or random algorithm creation (press 2).

### Subjective and objective shape evaluations

Following the scan, subjects evaluated each shape’s beauty level on a 1-4 scale (4 being most beautiful), chose the 20 weirdest shapes, and also chose the most iconic shapes in blocks of 20 shapes.

Block’s beauty score was calculated as the summation of beauty scores of all the shapes in the block. Weirdness and iconicity scores were calculated independently, in the same manner; A shape’s weirdness/iconicity score was equal to the number of subjects which rate the shape as weird/iconic. A block’s weirdness/iconicity score was calculated as the summation of weirdness/iconicity scores of all shapes in the block. In addition- the shapes were analyzed according to symmetry as an objective parameter. Block’s symmetry was calculated as the summation of all rotation and reflection symmetry groups of all the shapes in the block.

### MRI Data Acquisition and Preprocessing

The data were acquired on a 3 Tesla Trio Magnetom Siemens scanner at the Weizmann Institute of Science. Functional images of blood oxygenation level dependent (BOLD) contrast comprising of 46 axial slices were obtained with a T2*-weighted gradient echo planar imaging (EPI) sequence (3 × 3 × 3 mm voxel, TR = 3000 ms, TE = 30, flip angle = 75°, FOV 240 mm) covering the whole brain. Anatomical images for each subject were acquired in order to incorporate the functional data into the 3D Talairach space (Talairach and Tournoux, 1988) using 3-D T1-weighted images with high resolution (1 × 1 × 1 mm voxel, MPRAGE sequence, TR= 2300 ms, TE= 2.98 ms).

The first 7 images of each functional scan (including the extra initial block and rest) were discarded. Functional scan preprocessing included 3D motion correction and filtering out of low frequency noise (slow drift), and spatial smoothing using an isotropic Gaussian kernel of 6 mm full-width-half-maximum (FWHM). The functional images were superimposed on 2D anatomic images and incorporated into the 3D data sets through trilinear interpolation. Statistical analysis was based on a general linear model in which all stimuli conditions were defined as predictors, and convolved with the hemodynamic response function (HRF).

### Data analysis

In order to learn about the connection between shapes’ characteristics, perception and brain activity several measurements were examined; Reaction times (both tasks), response accuracy (*human*/*random*, task2), subjective evaluations post-scan (beauty, weirdness and iconicity), and symmetry score.

To investigate a possible difference in reaction times between shape categories (*random, not chosen, chosen, top rated*), Mann-Whitney tests were calculated for experiment 1 and 2 within subject and between subjects. Aesthetic (beauty) ratings for each shape were collected by each subject post the scan (on a 1-4 scale). To control for difference in rating patterns each subject’s ratings were Z normalized. Five GLM analyses were conducted; the first included four predictors: *random, not chosen, chosen, top rated*. A second GLM analysis with two predictors based on the category’s creator: *human* or *random*. In addition, in order to relate subjective blocks’ characteristics to brain activity, three parametric GLM analyses were conducted for beauty, weirdness and iconicity. In these multi-subject, random effect analyses each block of shapes received a weight according to its score (beauty/weirdness/iconicity) which was represented in the model as differential amplitude of the BOLD signal. Beauty, being an individual score calculated per subject separately, was z normalized between subjects.

Model selection of beauty and weirdness as well as beauty and iconicity fit to the probability to be classified as human was done based on the spearman correlation between model predictions and the Akaike information criterion (AIC). Models were generated as all possible combinations of the three parameters, either alone or coupled together. In order to have the monotonicity of the two scores increase in the same direction, weirdness score was represented as *e*^*−Weirdness score*^ (see SI for more details).

In all GLM analyses beta coefficients were calculated for the regressors, and a Student’s t-test was performed. Multi-subject analysis was based on a random-effect GLM. Multi-subject contrast maps (*human* vs. *random*, or category vs. category) were projected on an unfolded, inflated Talairach-normalized brain. Significance levels were calculated, taking into account the minimum cluster size and the probability threshold of a false detection of any given cluster. This was accomplished by a Monte Carlo simulation (cluster-level statistical threshold estimator in “Brain Voyager” software).

In the first experiment (task1) for *human* vs. *random* contrast (Fig. 3) a minimum cluster size of 103 voxels was significant. A minimum cluster size of 71 voxels was significant for *top rated* vs. *chosen*, 78 voxels for both *top rated* vs. *not chosen*, and *chosen* vs. *not chosen* (sup Fig. 2a). The minimum significant cluster size for each *human* category vs. *random* (sup Fig. 2b) was 103 voxels (*top rated*), 91 voxels (*chosen*) and 85 voxels (*not chosen*). For the parametric maps (Fig. 4) a minimum cluster size of 84 voxels was significant for beauty in task 1, and 90 voxels in task 2. For weirdness a minimum cluster size of 98 voxels was significant in task 1, and 105 voxels in task 2. For iconicity a minimum cluster size of 90 voxels was significant in task 1, and 104 voxels in task 2.

### ROI definition and analysis

Analysis of LOC-relevant voxels was conducted by defining a group bilateral ROI within the LOC using the contrast *human* > *random* in one task, and sampled in the other task. Note that the inverted contrast i.e. *random*> *human* failed to reveal any voxels in the LOC region (see Fig. 3). The ROI’s averaged beta weight (across voxels) was calculated per subject, for each predictor. Two-tailed paired t-tests (within subjects) were conducted between *human* and *random* beta weights for unaware (task 1) and aware (task 2) stimuli. A beta weight was extracted for each block and was plotted as a function of subjective beauty and symmetry (Fig. 5). Spearman correlation was calculated between participant’s average beta for each block (averaged over each ROI) and their aforementioned features.

## Acknowledgements

We thank Nahum Stern, Fanny Atar and Dr. Edna Haran-Furman for their assistance in the imaging setup and fMRI data collection. We thank Dr. Lior Noy for developing the shapes-creation computer game. This study was supported by the EU FP7 VERE, EU HBP Flagship (R.M.), ICORE ISF grants (R.M.), the Helen and Martin Kimmel award (R.M.), the Braginsky Center for the Interface Between Science and Humanities (U.A). U.A. is the incumbent of the Abisch-Frenkel Professorial Chair.

---

**Supplementary Figure 1.**
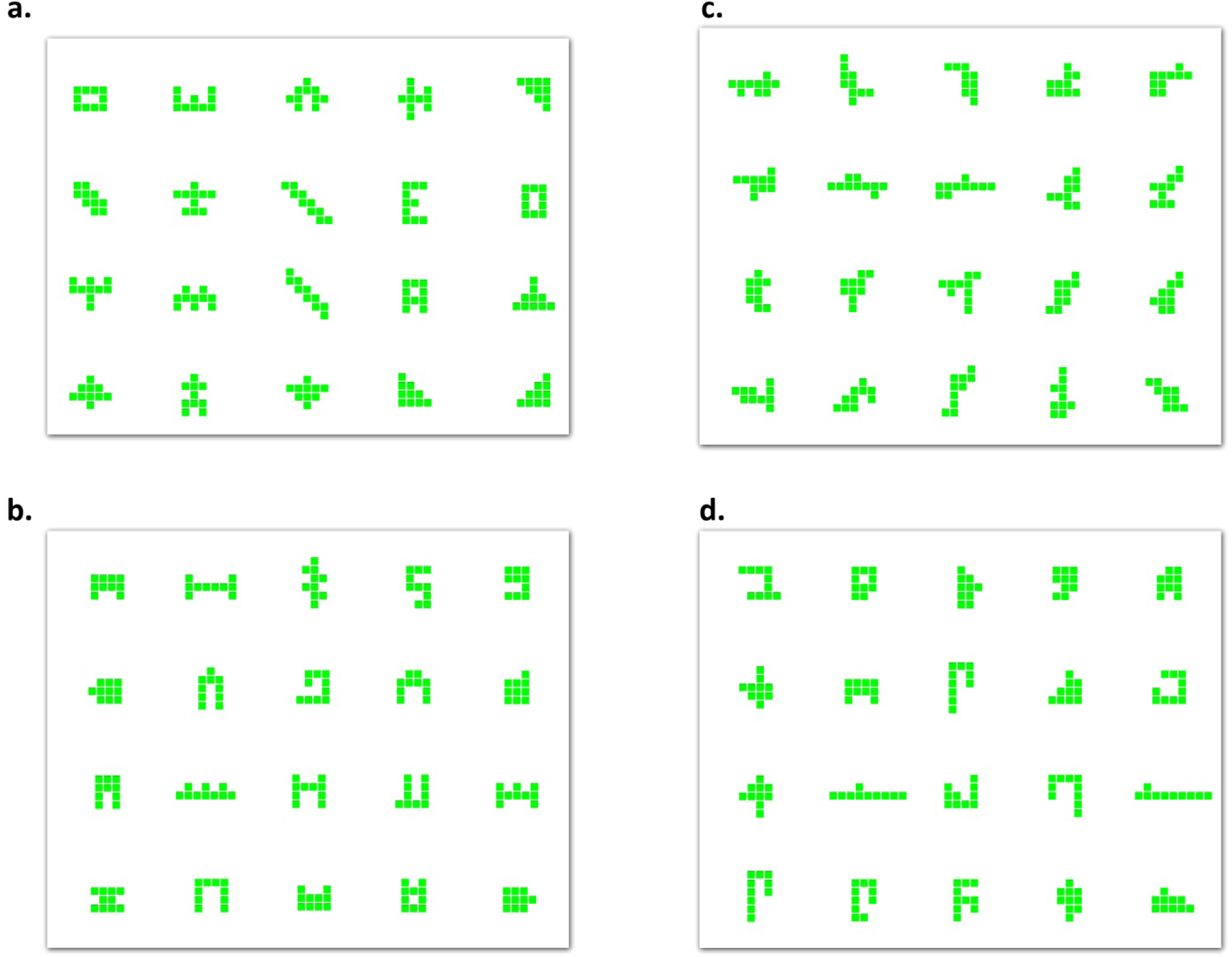
The entire stimuli ensemble. (a) *Top rated* shapes, (b) *Chosen* shapes, (c) *Random* - Never created shapes, (d) *Not chosen* shapes.

**Supplementary Figure 2.**
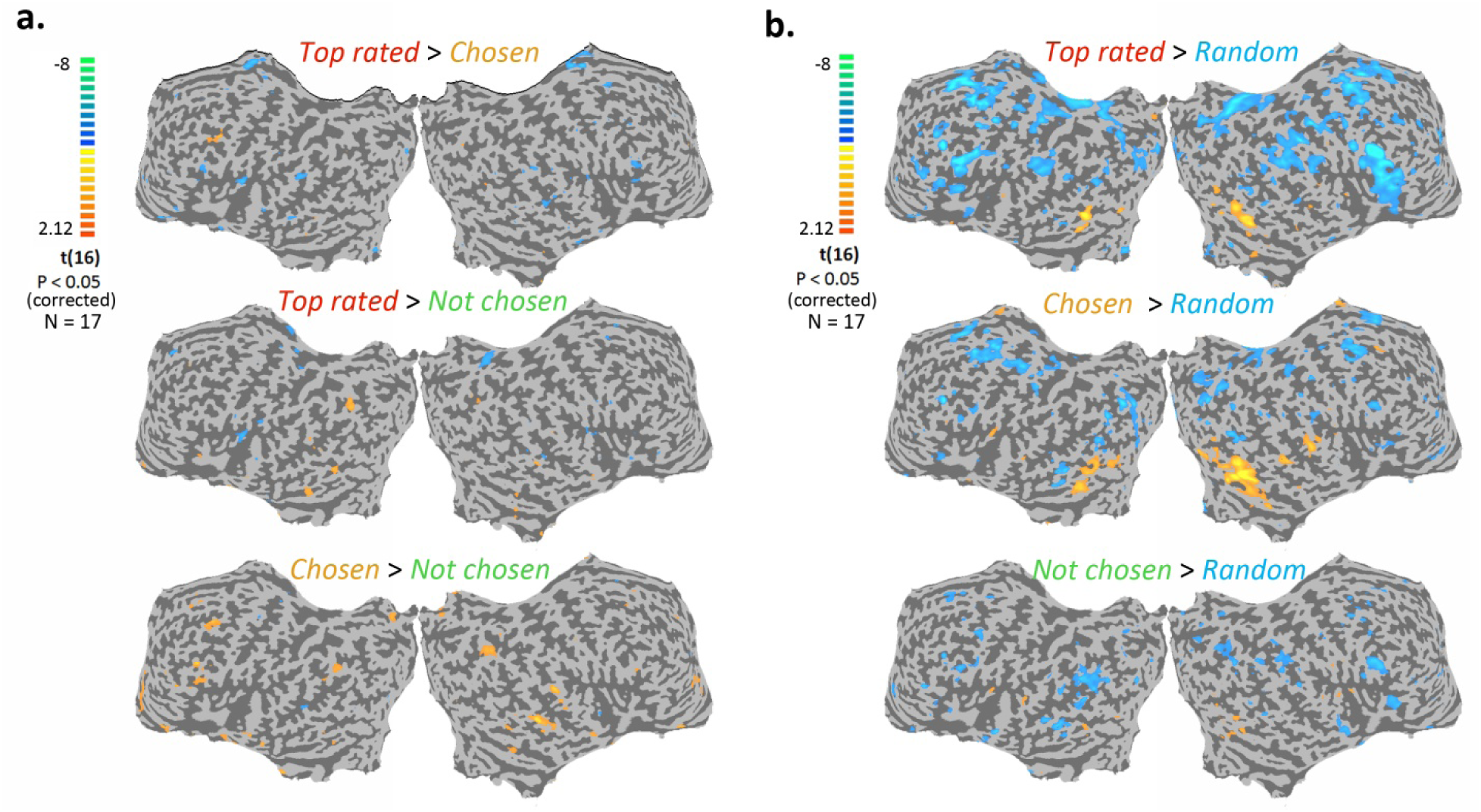
Comparison cortical activations between categories (experiment 1). Multi subjects activity maps (N = 17, corrected p < 0.05) are presented on an unfolded cortex. Color scale indicates t values. (a) Contrast maps between all the human generated shapes categories *(Human).* Yellow-orange scale represents regions which were more activated while watching blocks from the category in the left side of the contrast. Blue-green scale represents regions which were more activated while watching blocks from the category in the right side of the contrast. (b) Contrast maps between the *Human* categories and *Random* category (computer generated shapes). Color scale indicates t values. Yellow-orange scale represents regions which were more activated while watching shapes of *human* categories, compared to *random* shapes (blue-green scale).

## Supplementary Information

### Modelling shapes’ probability to be classified as human created using the behavioral scores of subjects

Here we describe the model selection process of finding the best model for the probabilities to be classified as human-created vs. random created shapes. The probability function was fit using a Logit function, 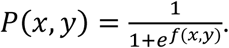 The variables of the model were the three subjective scores in our experiment – beauty (denoted as *b*), weirdness (denoted as *w*) and iconicity (denoted as *i*). Since weirdness and iconicity carry inverse correlations to the behavioral data (iconicity increases the probability to be classified as human, while weirdness decreases that same probability), we chose to map the weirdness score in the following way - *W*^’^ → *e*^−*w*^.

We tested all possible linear combinations of the individual scores, their pair and triplet interactions, yielding:

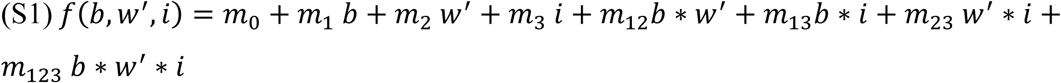

We assessed each model’s correlation with the data as well as its Akaike Information Criterion (AIC) score to attain the most accurate and simplest best fit model. The most accurate model with least number of parameters is one containing the interaction term between beauty and weirdness as a sole parameter, suggesting that it is the combination of beauty and familiarity that is the dominant component of classifying a shape as having human origin. In Table S1 we list the highest correlation and AIC scores of the different models for both beauty and weirdness and beauty and iconicity.

**Table S1:** Pearson correlation, p-values and AIC scores of Logit models with beauty, weirdness and iconicity scores.

|  | Model | Model parameters | Spearman correlation | P-value | AIC scores |
| --- | --- | --- | --- | --- | --- |
| 1 | $f(b, w', i) = b * w'$ | $\frac{1}{1 + e^{2.6 - 6.6 b w'}}$ | 0.96 | $4 * 10^{-21}$ | 5.84 |
| 2 | $f(b, w', i) = i$ | $\frac{1}{1 + e^{1.5 - 7 i}}$ | 0.94 | $5 * 10^{-19}$ | 6.32 |
| 3 | $f(b, w', i) = i * w'$ | $\frac{1}{1 + e^{1.4 - 7 i w'}}$ | 0.94 | $1 * 10^{-18}$ | 6.4 |
| 4 | $f(b, w', i) = b$ | $\frac{1}{1 + e^{3.8 - 8 b}}$ | 0.93 | $5 * 10^{-17}$ | 6.54 |
| 5 | $f(b, w', i) = b * i$ | $\frac{1}{1 + e^{1.3 - 9.2 b i}}$ | 0.92 | $5 * 10^{-16}$ | 7.04 |
| 6 | $f(b, w', i) = b * w' * i$ | $\frac{1}{1 + e^{1.2 - 9.3 b i w'}}$ | 0.92 | $1 * 10^{-15}$ | 7.13 |
| 7 | $f(b, w', i) = w'$ | $\frac{1}{1 + e^{7.1 - 9 w'}}$ | 0.89 | $2 * 10^{-12}$ | 5.36 |

## References

Aalto, S., N T Nen, P., Wallius, E., Mets Honkala, L., Stenman, H., Niemi, P. & Karlsson, H. 2002. Neuroanatomical substrata of amusement and sadness: a PET activation study using film stimuli. Neuroreport, 13, 67.

Beck, D. M., Pinsk, M. A. & Kastner, S. 2005. Symmetry perception in humans and macaques. Trends in cognitive sciences, 9, 405–406.

Brown, S., Gao, X., Tisdelle, L., Eickhoff, S. B. & Liotti, M. 2011. Naturalizing aesthetics: brain areas for aesthetic appraisal across sensory modalities. Neuroimage, 58, 250–258.

Cattaneo, Z., Lega, C., Ferrari, C., Vecchi, T., Cela-Conde, C. J., Silvanto, J. & Nadal, M. 2015. The role of the lateral occipital cortex in aesthetic appreciation of representational and abstract paintings: A TMS study. Brain and cognition, 95, 44–53.

Cela-Conde, C. J., Marty, G., Maestú, F., Ortiz, T., Munar, E., FernáNdez, A., Roca, M., Rosselló, J. & Quesney, F. 2004. Activation of the prefrontal cortex in the human visual aesthetic perception. Proceedings of the National Academy of Sciences of the United States of America, 101, 6321–6325.

Chatterjee, A., Thomas, A., Smith, S. E. & Aguirre, G. K. 2009. The neural response to facial attractiveness. Neuropsychology, 23, 135.

Chauvet, J.-M., Brunel Deschamps, E. & Hillaire, C. 1996. Dawn of art: the Chauvet Cave: the oldest known paintings in the world, HN Abrams.

De Beeck, H. P. O., Baker, C. I., Dicarlo, J. J. & Kanwisher, N. G. 2006. Discrimination training alters object representations in human extrastriate cortex. The Journal of Neuroscience, 26, 13025–13036.

Deshpande, G., Hu, X., Lacey, S., Stilla, R. & Sathian, K. 2010. Object familiarity modulates effective connectivity during haptic shape perception. Neuroimage, 49, 1991–2000.

Di Dio, C., Macaluso, E. & Rizzolatti, G. 2007. The golden beauty: brain response to classical and renaissance sculptures. PloS one, 2, e1201.

Fiez, J. A. & Petersen, S. E. 1998. Neuroimaging studies of word reading. Proceedings of the National Academy of Sciences, 95, 914–921.

Gauthier, I., Tarr, M. J., Anderson, A. W., Skudlarski, P. & Gore, J. C. 1999. Activation of the middle fusiform’face area’increases with expertise in recognizing novel objects. Nature neuroscience, 2, 568–573.

Grill-Spector, K., Kourtzi, Z. & Kanwisher, N. 2001. The lateral occipital complex and its role in object recognition. Vision research, 41, 1409–1422.

Hasson, U., Levy, I., Behrmann, M., Hendler, T. & Malach, R. 2002. Eccentricity bias as an organizing principle for human high-order object areas. Neuron, 34, 479–490.

Hodgson, D. 2009. Evolution of the visual cortex and the emergence of symmetry in the Acheulean techno-complex. Comptes Rendus Palevol, 8, 93–97.

Huettel, S. A. & Mccarthy, G. 2004. What is odd in the oddball task?: Prefrontal cortex is activated by dynamic changes in response strategy. Neuropsychologia, 42, 379–386.

Ishizu, T. & Zeki, S. 2011. Toward a brain-based theory of beauty. PloS one, 6, e21852.

Johansson, G. 1973. Visual perception of biological motion and a model for its analysis. Attention, Perception, & Psychophysics, 14, 201–211.

Kanwisher, N., Chun, M. M., Mcdermott, J. & Ledden, P. J. 1996. Functional imaging of human visual recognition. Cognitive Brain Research, 5, 55–67.

Kanwisher, N., Woods, R. P., Iacoboni, M. & Mazziotta, J. C. 1997. A locus in human extrastriate cortex for visual shape analysis. Cognitive Neuroscience, Journal of, 9, 133–142.

Kawabata, H. & Zeki, S. 2004. Neural correlates of beauty. Journal of Neurophysiology, 91, 1699–1705.

Kirk, U. 2008. The neural basis of object-context relationships on aesthetic judgment. PloS one, 3, e3754.

Lacey, S., Flueckiger, P., Stilla, R., Lava, M. & Sathian, K. 2010. Object familiarity modulates the relationship between visual object imagery and haptic shape perception. Neuroimage, 49, 1977–1990.

Lacey, S., Hagtvedt, H., Patrick, V. M., Anderson, A., Stilla, R., Deshpande, G., Hu, X., Sato, J. R., Reddy, S. & Sathian, K. 2011. Art for reward’s sake: Visual art recruits the ventral striatum. Neuroimage, 55, 420–433.

Lerner, Y., Hendler, T. & Malach, R. 2002. Object-completion effects in the human lateral occipital complex. Cerebral Cortex, 12, 163–177.

Malach, R., Reppas, J., Benson, R., Kwong, K., Jiang, H., Kennedy, W., Ledden, P., Brady, T., Rosen, B. & Tootell, R. 1995. Object-related activity revealed by functional magnetic resonance imaging in human occipital cortex. Proceedings of the National Academy of Sciences, 92, 8135–8139.

Mccandliss, B. D., Cohen, L. & Dehaene, S. 2003. The visual word form area: expertise for reading in the fusiform gyrus. Trends in cognitive sciences, 7, 293–299.

Murray, S. O., Kersten, D., Olshausen, B. A., Schrater, P. & Woods, D. L. 2002. Shape perception reduces activity in human primary visual cortex. Proceedings of the National Academy of Sciences, 99, 15164–15169.

Noy, L., Hart, Y., Andrew, N., Ramote, O., Mayo, A. & Alon, U. Year. A quantitative study of creative leaps. In: Proc. Third Int. Conf. Comput. Creativity, Dublin, 2012. Citeseer, 72–76.

Rotshtein, P., Malach, R., Hadar, U., Graif, M. & Hendler, T. 2001. Feeling or Features:: Different Sensitivity to Emotion in High-Order Visual Cortex and Amygdala. Neuron, 32, 747–757.

Sasaki, Y., Vanduffel, W., Knutsen, T., Tyler, C. & Tootell, R. 2005. Symmetry activates extrastriate visual cortex in human and nonhuman primates. Proceedings of the National Academy of Sciences of the United States of America, 102, 3159–3163.

Shibata, H., Inui, T. & Ogawa, K. 2011. Understanding interpersonal action coordination: an fMRI study. Experimental Brain Research, 211, 569–579.

Simion, F., Regolin, L. & Bulf, H. 2008. A predisposition for biological motion in the newborn baby. Proceedings of the National Academy of Sciences, 105, 809–813.

Talairach, J. & Tournoux, P. 1988. Co-planar stereotaxic atlas of the human brain. 3-Dimensional proportional system: an approach to cerebral imaging.

Tsao, D. Y., Freiwald, W. A., Knutsen, T. A., Mandeville, J. B. & Tootell, R. B. 2003. Faces and objects in macaque cerebral cortex. Nature neuroscience, 6, 989–995.

Vartanian, O. & Goel, V. 2004. Neuroanatomical correlates of aesthetic preference for paintings. Neuroreport, 15, 893–897.

